# Unbiased discovery of antibody therapies that stimulate macrophage-mediated destruction of B-cell lymphoma

**DOI:** 10.1101/2024.11.13.623229

**Authors:** Juliano Ribeiro, Carlota Pagès-Geli, Anna Meglan, Jose Velarde, Jasmine Blandin, Kyle Vaccaro, Thomas Wienclaw, Patricia Fernández-Guzmán, Cynthia K. Hahn, Marta Crespo, Kipp Weiskopf

## Abstract

Macrophages are critical effectors of antibody therapies for lymphoma, but the best targets for this purpose remain unknown. Here, we sought to define a comprehensive repertoire of cell surface antigens that can be targeted to stimulate macrophage-mediated destruction of B-cell lymphoma. We developed a high-throughput assay to screen hundreds of antibodies for their ability to provoke macrophages to attack B-cell lymphoma cells. Across both mouse and human systems, we identified multiple unappreciated targets of opsonization as well as putative immune checkpoints. We used this information to engineer a compendium of 156 bispecific antibodies, and we identified dozens of bispecifics that dramatically stimulate macrophage-mediated cytotoxicity of lymphoma cells. Among these, a bispecific comprising a SIRPα decoy domain and a CD38-targeting arm (WTa2d1xCD38) exhibited maximal efficacy while minimizing the risk of hematologic toxicity. This bispecific stimulated robust anti-tumor responses in multiple xenograft models of aggressive B-cell lymphoma. Our approach can be directly applied to other cancers to rapidly discover bispecific antibodies that leverage anti-tumor responses by macrophages or other innate immune cells.

---

B-cell non-Hodgkin lymphoma (NHL) is the most common hematologic malignancy worldwide and remains with a large, unmet clinical need. It is a highly heterogeneous disease, with multiple subtypes distinguished by variations in both genetic alterations and the immune microenvironment^1,2^. Diffuse large B-cell lymphoma (DLBCL) is the most common subtype and accounts for approximately 35% of all B-cell lymphomas. Despite advances in frontline rituximab-based chemoimmunotherapy (e.g., R-CHOP), a significant proportion of patients become refractory or relapse after the first line of treatment^3^. Subsequent treatment strategies have focused on promoting T-cell cytotoxicity, such as CAR T-cell therapy and bispecific T cell engagers. However, these therapies only cure a subset of patients^4,5^, and many patients exhibit inadequate T-cell infiltration and function^6–8^. Additionally, more than 50% of B-cell lymphomas exhibit genetic alterations that cause these cells to evade recognition by T cells, such as downregulation or loss of the Major Histocompatibility Complex (MHC) antigen presentation machinery or overexpression of immune checkpoint receptors^9^. While these lymphomas may be less vulnerable to T-cell-dependent killing, these features create opportunities for orthogonal approaches that target innate immune cells, particularly macrophages.

In B-cell lymphoma, the presence of CD68+ macrophages was linked to favorable prognosis in patients treated with R-CHOP, while it correlated with poor outcomes in the absence of rituximab^10^. Antibody-dependent cellular phagocytosis (ADCP) is a major mechanism that contributes to the efficacy of rituximab in vivo^11^, likely underlying this effect. However, ADCP is often limited by the CD47/SIRPα macrophage immune checkpoint. CD47 is a cell-surface antigen expressed by many normal and neoplastic tissues and acts as a “don’t eat me” signal to prevent macrophage phagocytosis^12^. Therapies that block the CD47/SIRPα interaction can enhance ADCP and synergize with rituximab in B-cell lymphoma preclinical models^13–15^. This therapeutic approach has been extended to clinical trials and shows promising efficacy in patients with B-cell malignancies^16^. Similarly, downregulation of MHC-I may make lymphoma cells more vulnerable to macrophage phagocytosis in the presence of tumor-opsonizing antibodies or CD47-blocking therapies^17^.

Despite the success of monoclonal antibodies for hematologic malignancies, there have been limited efforts to identify additional antibodies that stimulate macrophages to attack lymphoma cells. To address this limitation, we developed a high-throughput system to screen libraries of monoclonal antibodies to a diversity of cell surface antigens. Our approach is target agnostic and “function first” to identify and prioritize antibodies that robustly stimulate macrophages to eliminate lymphoma cells. We successfully performed screens across both the murine and human systems, and we used the results from our screens to engineer an extensive collection of bispecific antibodies (bsAbs) that could maximally activate macrophage anti-tumor functions against aggressive B-cell lymphomas. Overall, we have identified therapeutic strategies and therapeutic agents to provoke macrophages to attack B-cell lymphoma. These efforts can be translated to the clinic, and we expect this approach can be rapidly applied to identify new therapies for other types of cancer.

## Results

### Development of a high-throughput system to screen for macrophage-mediated cytotoxicity of B-cell lymphoma

To identify unappreciated targets for macrophage-directed immunotherapy of B-cell lymphoma, we adopted a high-throughput assay^18^ to screen antibodies for their ability to stimulate macrophage-mediated cytotoxicity of lymphoma cells (**Fig. 1a**). In our initial approach, we co-cultured mouse M2-like macrophages^19–21^ with fluorescent A20 MHC-I KO lymphoma cells. We employed an MHC-I KO line to model intrinsic resistance to T cell-directed killing^22–24^ and focus on opportunities for innate immune intervention. To the co-cultures we added an arrayed library of 173 purified monoclonal antibodies (**Supplementary Table 1**) targeting different murine cell-surface antigens. We tested each antibody in duplicate across three different treatment conditions: (i) monotherapy (antibodies alone), (ii) combination with anti-CD47, or (iii) combination with anti-CD20. We performed automated microscopy and image analysis over a 7-day period to quantify the fluorescent area as a metric of lymphoma cell growth or elimination. For each treatment condition, we identified the antibodies that stimulated the greatest anti-tumor function. Consistently across all conditions, we observed that antibodies to CD24, CD95 (FAS), CD40, CD184 (CXCR4), and CD79b were able to elicit robust macrophage-mediated cytotoxicity of the A20 cells (**Fig. 1b,c,d,e**). We also performed quantitative surfaceome profiling by evaluating the ability of each antibody to bind to the surface of the A20 cells by flow cytometry. We observed a modest correlation between antibody binding and the anti-tumor response by macrophages (**Fig. 1f,g,h**). For example, CD24 was the highest detected surface antigen in our assay and it induced the greatest anti-tumor response. However, many targets existed as outliers, suggesting that the degree of binding to lymphoma cells alone is not sufficient to predict whether an antibody can elicit macrophage-mediated cytotoxicity. Furthermore, our results indicate that some antibodies may preferentially act on the macrophages in the co-culture system (e.g., Ly-6A/E, 4-1BBL), suggesting they modulate immune checkpoints **(Supplementary Fig. 1)**. In total, these studies identified several unappreciated targets for innate immunotherapy on A20 cells that may present new therapeutic opportunities for lymphoma.

**Fig. 1.**
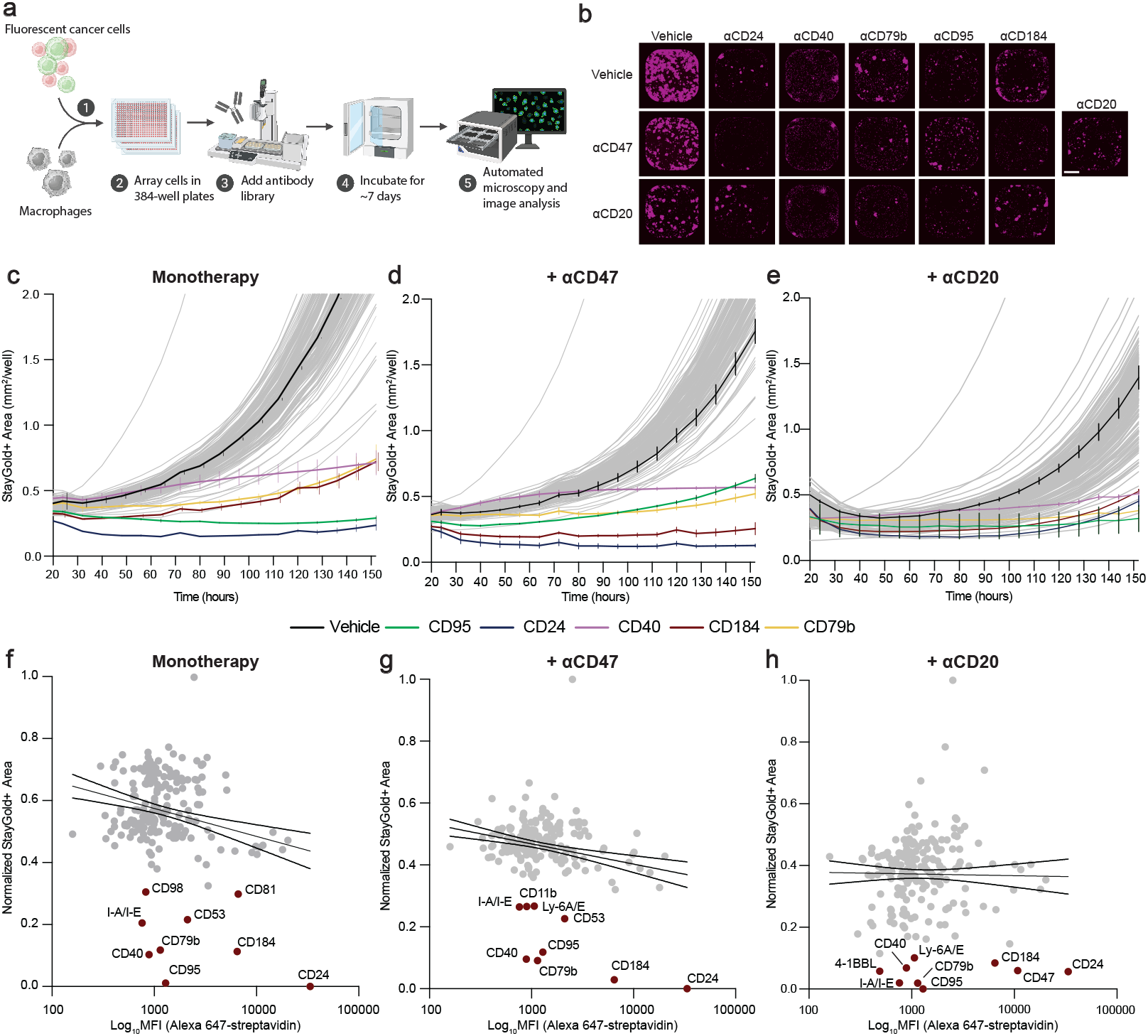
An unbiased antibody screen identifies targets for macrophage-directed immunotherapy of murine B-cell lymphoma. **a**, Experimental design of an unbiased functional screen to identify monoclonal antibodies that provoke macrophage-mediated cytotoxicity of lymphoma cells. Primary murine macrophages were co-cultured in 384-well plates with StayGold+ MHC-I A20 cancer cells (a murine B-cell lymphoma cell line). The wells were subjected to three treatment conditions: control (“monotherapy”), 10 µg/mL anti-CD47 antibody, or 10 ug/mL anti-CD20 antibody. An arrayed library of purified monoclonal antibodies targeting murine cell surface antigens (n = 173 antibodies) was overlaid at a concentration of 6.55 µg/mL. The cells were incubated for 156 h (∼6.5 days) and the StayGold+ area was quantified by automated microscopy and whole-well image analysis every 8 hours. **b**, Representative whole-well images at the last time point (156 h) showing StayGold+ lymphoma area (purple) from wells treated with the indicated antibodies that were found to stimulate macrophage-dependent cytotoxicity of A20 cells. Scale bar, 800 um. (**c-e**) Results of co-cultures with the antibody library showing growth of StayGold+ A20 MHC-I KO cells under monotherapy conditions (**c**), in combination with anti-CD47 (**d**), or in combination with anti-CD20 (**e**). Each curve represents the growth of lymphoma cells treated with a different antibody in the presence of macrophages. Black curve indicates control wells, colored curves indicate antibodies against CD24, CD40, CD79b, CD95 and CXCR4. Curves represent mean (± SEM as indicated) of two individual co-culture wells. **f-h**, Scatter plots depicting results of quantitative surfaceome profiling by flow cytometry for each library antibody binding to A20 lymphoma cells (x-axis) versus anti-lymphoma function at the last imaging time point (t = 156 hours, y-axis). Lower fluorescent area values indicate greater macrophage anti-lymphoma activity. Highlighted in red are the antibodies exceeding the 95^th^ percentile in functional activity and are defined as hits. Black curve indicates the linear relationship with 95% CI boundaries between the functional efficacy and binding in the monotherapy condition (**f**, r=-0.2334, p=0.0011), in combination with anti-CD47(**g**, r=-0.2339, p=0.0011), and in combination with anti-CD20 (**h**, r=-0.009, p=0.8987).

### Discovery of targets and antibodies for macrophage-mediated cytotoxicity of human B-cell lymphoma

To understand whether we could similarly identify targets of antibody-dependent phagocytosis in the human system, we also evaluated an arrayed library of 241 purified monoclonal antibodies (**Supplementary Table 2**) targeting different human cell-surface antigens (**Fig. 2**). The library included antibodies to 92 antigens that were shared with the murine library (**Supplementary Fig. 2**). We used primary human macrophages and co-cultured them with GFP+ Raji cells, an aggressive Burkitt lymphoma cell line. As with the murine system, we tested all of the antibodies in three conditions: (i) monotherapy (antibodies alone), (ii) combination with anti-CD47, and (iii) combination with anti-CD20 (**Fig. 2a,b,c**). We identified functional antibodies that were consistent across all three treatment conditions. Of note, a strong association was observed with antibodies targeting MHC molecules on the lymphoma cells, including antibodies to MHC-I (HLA A,B,C, β2M), and MHC-II (HLA-DR-DQ-DP, HLA-DR) components. Moreover, we identified CD71, CD184, CD45, CD98, CD38, and CD147 blocking antibodies as top hits across multiple treatment conditions. Again, we observed a modest correlation between the degree of antibody binding and elimination of the lymphoma cells in culture with some notable standouts (**Fig. 2d,e,f**). As an example, an anti-CD85j (LILRB1) antibody had no substantial effect as a single agent, but exhibited the greatest anti-tumor response in combination with anti-CD47 (**Fig. 2d,e**). This observation is consistent with the reported function of CD85j (LILRB1) as a macrophage immune checkpoint^17^, and indicates this screening platform can be useful to identify antibodies that act by either opsonizing the lymphoma cells, exerting a functional effect directly on the macrophages, and/or cross-linking macrophages and lymphoma cells to promote phagocytosis.

**Fig. 2.**
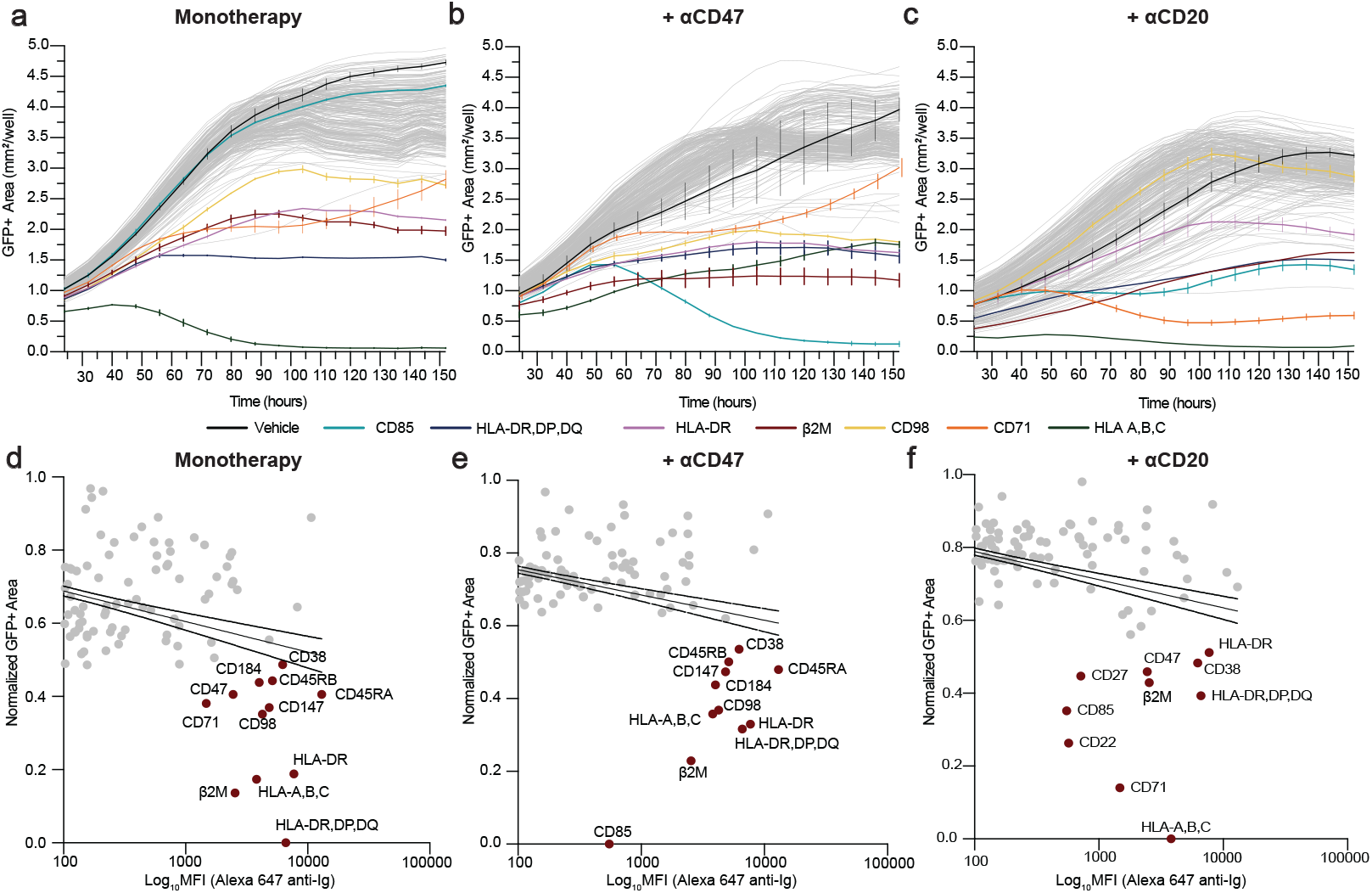
An unbiased antibody screen identifies targets for macrophage-directed immunotherapy for human B-cell lymphoma. Primary human macrophages were co-cultured in 384-well plates with GFP+ Raji cells (a human Burkitt lymphoma cell line). The wells were subjected to three treatment conditions: control, 10 µg/mL anti-CD47 antibody, or 10 µg/mL anti-CD20 antibody. An arrayed library of purified monoclonal antibodies targeting human cell surface antigens (n = 241 antibodies) was overlaid at a concentration of 6.55 µg/mL. The cells were then incubated for 156 h (∼6.5 days) and the GFP+ area was quantified by automated microscopy and whole-well image analysis every 8 hours. **a-c**, Growth of GFP+ Raji cells in co-culture with macrophages under monotherapy conditions (**a**), in combination with anti-CD47 (**b**), or in combination with anti-CD20 (**c**). Each curve represents the growth of lymphoma cells treated with a different antibody. Black curve indicates control wells, colored curves indicate top antibody hits against MHC-related antigens, CD85, CD98 and CD71. Curves represent mean (± SEM as indicated) of two individual co-culture wells. **d-f**, Scatter plots depicting results of quantitative surfaceome profiling by flow cytometry for each library antibody binding to GFP+ Raji lymphoma cells (x-axis) versus functional anti-lymphoma effectiveness at the last imaging time point (t = 156 hours, y-axis). Lower fluorescent area values indicate greater macrophage anti-lymphoma activity. Highlighted in red are the antibodies exceeding the 95^th^ percentile in function activity and are defined as hits. Black curve indicates the linear relationship with 95% CI boundaries between the functional efficacy and binding in the monotherapy condition (**d**, r=-0.3364, p<0.0001), in combination with anti-CD47 (**e**, r=-0.3646, p<0.0001), and in combination with anti-CD20 (**f**, r=-0.4105, p<0.0001).

### Antibody combinations to maximize macrophage-mediated cytotoxicity of B-cell lymphoma

Together, our murine and human studies identified several therapeutic targets and corresponding antibodies that could stimulate macrophage-mediated cytotoxicity of lymphoma cells. Since combinations of therapeutic antibodies have shown promise in ongoing clinical trials for patients with lymphoma, particularly combinations of anti-CD47 and anti-CD20 antibodies^16^, we next evaluated whether combinations of antibodies to the targets we identified could elicit more robust anti-tumor responses in vitro in the murine and human systems. Given the central importance of MHC-I molecules in regulating macrophage phagocytosis, we examined both wild-type and MHC-I knockout cell lines. Using syngeneic mouse macrophages and A20 lymphoma cells, we evaluated the combination of the top five antibodies from the murine screen—anti-CD24, CD95 (FAS), CD40, CD184 (CXCR4), and CD79b—along with antibodies against CD47, CD20, and PD1 as comparisons. We individually combined all of these for a total of 64 different antibody combinations to identify those that exhibit the greatest activity for stimulating macrophage anti-lymphoma functions (**Fig. 3a,b,c**). We found that the A20 MHC-I knockout line was more vulnerable to ADCP but otherwise similar trends were observed across the wild-type and MHC-I knockout lines. Additionally, we found that there were several antibody combinations that were substantially more effective than even the combination of anti-CD20 and anti-CD47. In particular, most combinations with anti-CD24 or anti-CD95 antibodies caused elimination of nearly all lymphoma cells from the co-cultures (**Fig. 3a,b,c**). These findings suggest highly active antibody combinations can be identified that maximize the ability of macrophages to attack and eliminate lymphoma cells.

**Fig. 3.**
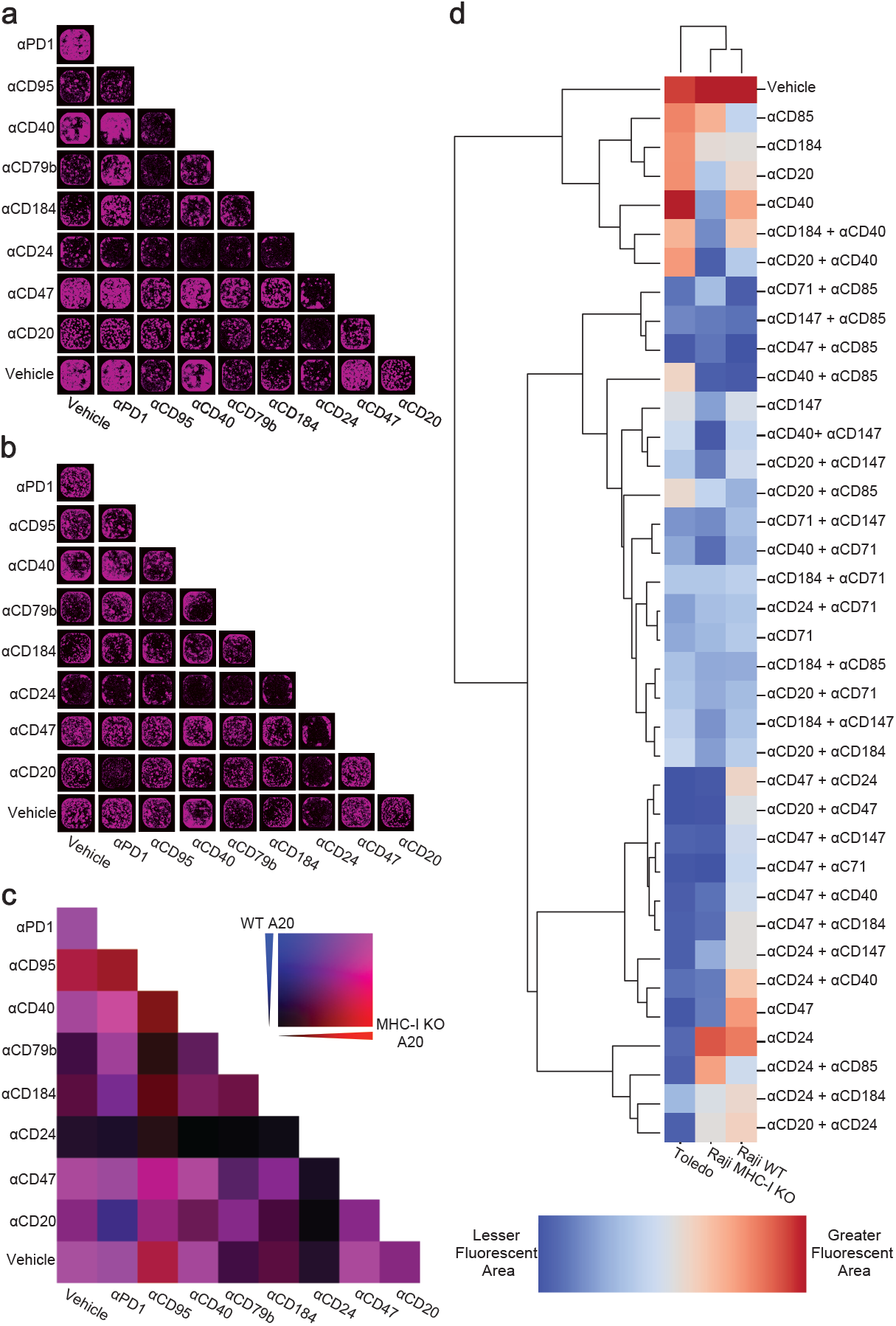
Identification of antibody combinations that elicit maximal macrophage-mediated cytotoxicity of B-cell lymphoma. **a,b**, Primary murine macrophages were co-cultured with StayGold+ A20 (WT) or MHC-I KO cells. Seven different antibodies identified from our screen and anti-PD1 were tested alone or in combination as indicated. GFP+ area was measured over time as a representation of growth or elimination of lymphoma cells. Representative images at the last time point (156 h) are shown for each antibody combination for StayGold+ A20 WT (**a**) and StayGold+ A20 MHC-I KO cells (**b**). **c**, Heat map representing StayGold+ area at last time point (152 h) for WT (blue) and MHC-I-negative (red) A20 cells. Merged data is shown in purple. For both cell lines, lighter color indicates greater StayGold+ area while darker color indicates lesser StayGold+ area. Data represents the mean of 2 individual co-cultures per antibody combination. **d**, Antibody combination studies were performed using primary human macrophages and GFP+ Raji (WT), GFP+ Raji MHC-I KO, and mScarlet+ Toledo cells treated with 8 different antibodies alone or in combination as indicated. Heatmap depicts the mean normalized fluorescent area of the last time point (152 h) from 2 individual co-culture wells. Unbiased hierarchical clustering was used to identify antibodies, combinations and cell lines with similar anti-lymphoma properties when co-cultured with macrophages.

Similarly, we evaluated eight antibody combinations in the human system. We tested wild-type Raji cells, Raji MHC-I KO cells, and Toledo cells (a human DLBCL cell line). The combinations included the top antibody hits from the human screen (anti-CD85, CD71, and CD147), the mouse system (anti-CD24 and CD40), and a shared antibody (anti-CD184), along with CD47 and CD20-targeting antibodies. From these efforts, we identified over a dozen antibody combinations that were highly active across all three cell lines tested. Among the most effective of these combinations was anti-CD85j (LILRB1) combined with either anti-CD47, anti-CD147, or anti-CD71 (**Fig. 3d**). Together, these findings indicate that highly active combination therapies can be identified, and the effectiveness of these combinations may be comparable to or exceed that of anti-CD20 combinations. Furthermore, the MHC-I knockout line was generally more sensitive to macrophage-dependent killing in response to antibodies, again underscoring the importance of MHC-I molecules in protecting lymphoma cells from macrophage phagocytosis.

### Development of a rapid system to create and evaluate bispecific antibodies for macrophage-mediated cytotoxicity

Our unbiased screening efforts and combination studies indicate that additional targets exist on lymphoma cells that can be leveraged to stimulate robust anti-tumor responses by macrophages. Some of these molecules may act as macrophage immune checkpoints, while others may serve as optimal targets of ADCP. Regardless of the mechanism of action, we reasoned that antibodies to the targets identified from our mouse and human screens could be adapted to a bispecific format to maximize anti-lymphoma responses by macrophages and generate therapies that would exhibit robust single-agent activity against lymphoma cells. We therefore developed a strategy to rapidly generate a total 156 bispecific macrophage-activating antibodies and screen them for the ability to stimulate macrophage-mediated cytotoxicity of lymphoma cells. To begin, we curated a library of available antibody sequences to targets identified from our screens (**Fig. 4a**). We formatted these into single-chain fragment variable constructs (scFvs), then fused them to a human IgG1 knob or hole Fc construct^25^. This heterodimeric scFv-Fc format permits the rapid combinatorial production and screening of bispecifics since each binding arm is encoded by a single chain that can be readily combined with other binding elements. Based on the results of our screening assays, we targeted CD47, CD24, CD79b, CD184, CD40, CD95, CD38, CD71, CD85j (LILRB1), and CD20. For targeting CD47, we tested two different single-domain binding modules: CV1 (a high-affinity SIRPα decoy protein), and WTa2d1 (a low affinity SIRPα decoy protein)^14^. We also targeted PD1 as a comparison. We crossed all of these binding arms with each other in two reciprocal formats (i.e., knob-hole and hole-knob), expressed them in Expi293F cells, and then evaluated the resulting antibodies for expression, binding, and function. We tested each bsAb in co-culture assays using primary human macrophages and three different human B-cell lymphoma cell lines: Raji, Toledo, and SUDHL-8 (**Fig. 4b,c,d,e,i**). We also examined the expression of each bsAb by ELISA and their binding to each of the cell lines **(Fig. 4f)**. Furthermore, since on-target red blood cell toxicity has been observed with some anti-CD47 antibodies in clinical trials^16,26^, we also examined the ability of each bispecific to bind to human red blood cells (**Fig. 4g**).

**Fig. 4.**
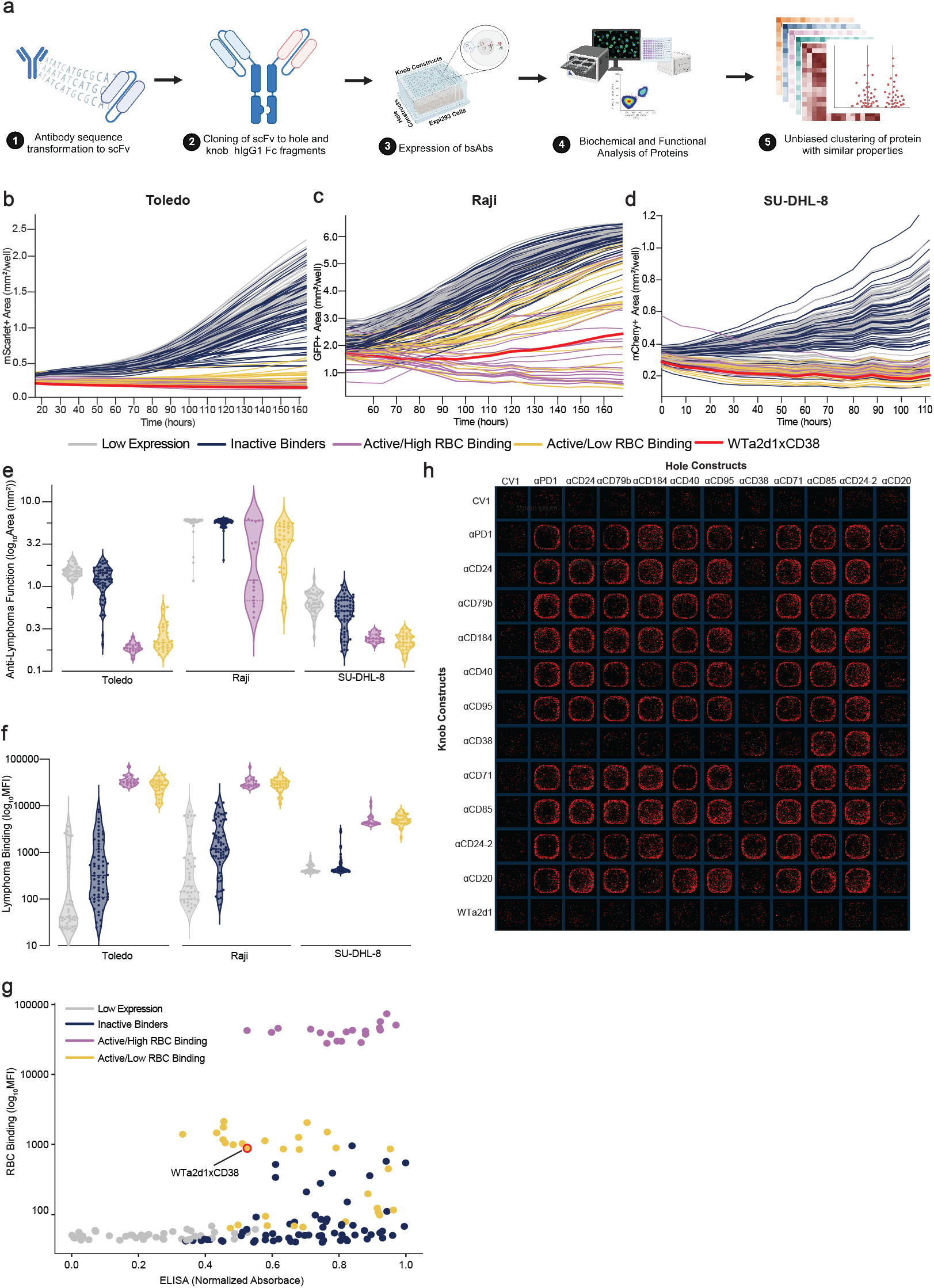
Multiplex generation of bispecific antibodies that stimulate macrophage destruction of human B-cell lymphoma. **a**, Experimental setup for the generation and functional characterization of a combinatorial matrix of bsAbs. scFvs were created based on publicly available antibody sequences, fused to modified human IgG1 Fc regions containing knob or hole mutations for bispecific assembly, and a combinatorial library of bsAbs was expressed in Expi293 cells. The resulting bsAbs were tested for functional activity and binding to multiple lymphoma cell lines and red blood cells. ELISA was performed to evaluate antibody expression. Finally, the properties of macrophage-mediated cytotoxicity, binding, and expression by ELISA were used for K-Means unbiased clustering of the antibodies with similar biochemical and functional properties. **b-d**, Growth of human lymphoma cells (**b**, mScarlet+ Toledo; **c**, GFP+ Raji; **d**, mCherry+ SUD-HL-8) in co-culture with primary human macrophages. Each curve depicts lymphoma growth in the presence of a different bsAb. Curves represent mean of 2 independent co-culture wells. Each color represents a different cluster of proteins based on functional properties. **e**, Violin plot of fluorescent area at the last time point for each lymphoma cell line for each unbiased cluster of bsAbs. **f**, Violin plot depicting binding of the antibodies to each lymphoma cell line as indicated based on clustering. **g**, Relationship between the different bsAb clusters with respect to binding to human red blood cells and their expression by ELISA. **h**, Representative images of the last time point of the co-culture of macrophages and mScarlet+ Toledo cells and each bsAb from the combinatorial library. Rows indicate hole constructs, columns indicate knob constructs. **e-f**, Violin plots depict mean and interquartile range.

Across all of these assays, we found that four distinct categories of bispecific emerged from unsupervised clustering analysis: (i) Bispecifics that were highly active and exhibited minimal red blood cell binding (‘Active/Low RBC binding’), (ii) Bispecifics that were highly active and exhibited intense red blood cell binding (‘Active/High RBC binding’), (iii) Bispecifics with limited activity, variable lymphoma binding, and low or absent red blood cell binding (‘Inactive binders’), and (iv) Bispecifics that did not fold or express well (‘Low Expression’). Consistently, we found that bispecifics formulated with a CV1 binding arm exhibited exceptional anti-tumor effects but also exhibited strong binding to red blood cells. Many bispecifics formulated with WTa2d1, CD20, or CD38 binding arms exhibited desirable properties of robust anti-lymphoma activity with minimal red blood cell binding. Other binding targets exhibited limited functional activity or did not express well (**Fig. 4e,f,g)**. Across all three cell lines tested, we found that a WTa2d1xCD38 bispecific was consistently among the most effective agents for stimulating macrophage-mediated cytotoxicity of B-cell lymphoma cells and exhibited minimal red blood cell binding (**Fig. 4b,c,d,g)**. Overall, these properties indicate this bispecific agent may be an ideal therapeutic for stimulating innate immune cells to attack and eliminate B-cell lymphoma.

### A WTa2d1xCD38 bispecific antibody is an optimal therapeutic candidate for B-cell lymphoma

Given the favorable properties observed using the WTa2d1xCD38 bispecific, we further evaluated the biochemical and functional qualities of this agent. We found that the predicted structure of this bispecific included an expected association via the T22Y(knob) and Y86T(hole) mutations (**Fig. 5a**) as previously described^25^. Moreover, by mass spectrometry, approximately 94% of the protein assembled in the expected bispecific format (**Fig. 5b**). In on-cell binding assays, we found that each arm is able to bind to its respective antigen **(Fig. 5c)** and that WTa2d1xCD38 binds to Raji, Toledo and human macrophages cells with an EC_50_ of 11.03, 0.37, and 2.90 nM, respectively (**Fig. 5d**), indicating high-affinity binding to lymphoma cells and macrophages with potential to bridge the two cell types. In macrophage co-culture assays using nine different human B-cell lymphoma cell lines, we found that the bsAb exhibited significant macrophage-mediated cytotoxicity with an IC_50_ ranging from 73 pM (SUDHL-8) to 3.17 nM (HBL-1) (**Fig. 5e**). For nearly all cell lines tested, the WTa2d1xCD38 bispecific was more potent and more effective than rituximab, a gold-standard clinical benchmark (**Fig. 5 f,g**). In phagocytosis assays, the bispecific elicited higher ADCP than both rituximab and anti-CD47 antibodies (**Fig. 5 h,i**). Finally, we assessed the contributions of the Fc gamma receptors (FcγR) in mediating responses to WTa2d1xCD38. We found that CD16-, CD32-, or CD64-blocking antibodies were able to decrease the activity of the bsAb in co-culture assays with macrophages, but the degree of inhibition varied based on the B-cell lymphoma cell line and the specific Fc receptor (**Fig. 5j**). These findings suggest the human IgG1 isotype is important for the effects of WTa2d1xCD38, but also that Fc-independent effects may contribute to its mechanism of action.

**Fig. 5.**
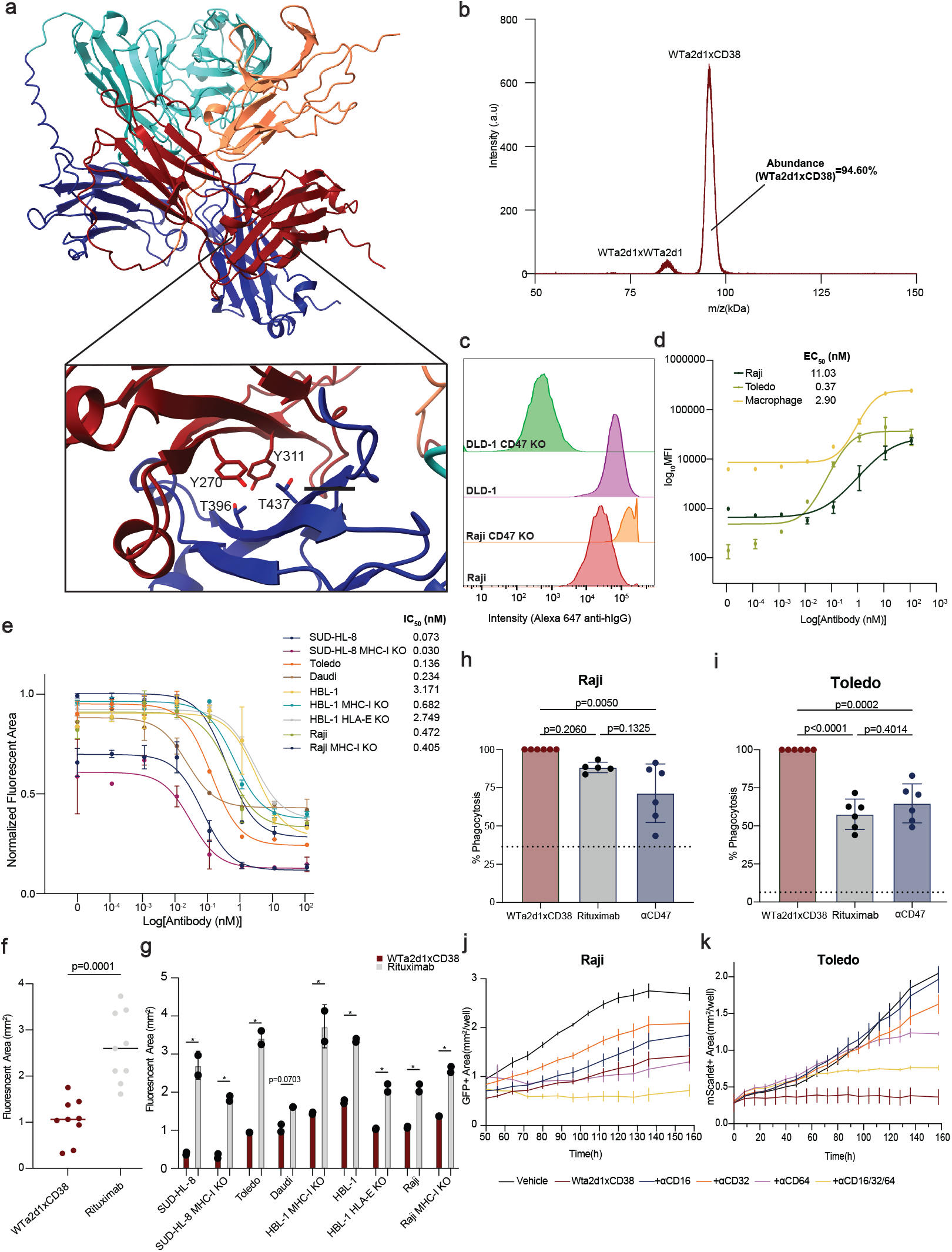
WTa2d1xCD38 stimulates maximal macrophage-mediated cytotoxicity of B-cell lymphomas. **a**, AlphaFold2^35^ predicted structure of WTa2d1xCD38, a bsAb targeting CD47 and CD38. The Fc region of the anti-CD38 arm is highlighted in blue with its respective scFv region in teal, whereas the Fc region of the WTa2d1 arm is highlighted in red with its respective binding domain in orange. The mutations, “knob” (T270Y,T311Y) and “hole” (Y396T, Y437T) are highlighted at the bottom. **b**, Maldi-TOF analysis of the protein population of the purified bsAb. Based on the theoretical mass of the possible homo and heterodimers, the two main peaks were assigned to WTa2d1xWTa2d1 and WTa2d1xCD38 species. The abundance of WTa2d1xCD38 was calculated as 94.60% based on the peak area. **c**, Histogram showing binding of the bsAb to wild-type versus CD47 knockout (KO) cell lines highlights the ability of the two antibody arms to bind to their respective antigens. Binding was detected with an Alexa 647-conjugated anti-human IgG secondary antibody. DLD-1 is a colorectal cancer cell line that expresses CD47 but not CD38. **d**, Binding of WTa2d1xCD38 to the indicated human lymphoma cell lines and macrophages combined from 3 donors indicates the cell-based affinity for the antibody. Data indicate mean ± SD. **e**, Co-culture assays using primary human macrophages were used to determine the potency of WTa2d1xCD38 across nine different human B-cell lymphoma cell lines, including the indicated wild-type and MHC-I KO variants. Each curve represents an 8-point titration performed in duplicate for each cell line. Fluorescent area was compared at the last imaging time point and normalized based on the maximum value for each cell line. Data depict mean ± SD. **f**, Comparison of the fluorescent area at the last time point for different B-cell lymphoma cell lines when treated with 10.54 µg/mL of WTa2d1xCD38 or rituximab. Each point represents the mean value for a different cell line performed in duplicate. Student two tailed paired T-test was used to evaluate the difference between the two groups. Lines show median. **g**, Comparison of the fluorescent area at the last time point from co-culture assays using the indicated lymphoma cell lines treated with either WTa2d1xCD38 or rituximab. Data depict mean ± SD. *p-value<0.0001 for the indicated comparisons by two-way ANOVA with correction for multiple comparisons. **h,i**, Macrophage phagocytosis of Raji (**h**) and Toledo (**i**) cells when co-cultured for 2 hours in the presence of the WTa2d1xCD38, rituximab, or an anti-CD47 antibody (clone B6H12). All of the values were normalized against the bsAb and the experiment was performed two independent times with three individual co-culture wells per condition. The dotted lines indicate the mean of the negative control condition performed in one experiment with three co-culture wells. Data depict mean ± SD with analysis by two way ANOVA with Tukey’s multiple comparison test. **j,k**, Co-culture with human macrophages pooled from multiple donors with Raji (**j**) and Toledo (**k**) lymphoma cells as targets. Co-cultures were treated with 1 μg/ml WTa2d1xCD38 and 10 μg/ml antibodies to different Fc gamma receptors as indicated. Curves indicate mean ± SEM.

To understand whether WTa2d1xCD38 could also exhibit therapeutic effects in vivo, we evaluated its efficacy in two xenograft models of aggressive B-cell lymphoma in NSG mice. These mice lack adaptive immune cells but contain myeloid cells capable of responding to macrophage-directed therapies^27^. First, we engrafted Raji cells subcutaneously into the flanks of NSG mice (**Fig. 6a**). After establishing tumors, we randomized the mice into three treatment groups: vehicle control, 100 µg WTa2d1xCD38, or 100 µg rituximab as a standard-of-care benchmark. We found that tumor growth was dramatically inhibited by treatment with either WTa2d1xCD38 or rituximab, compared to the vehicle control group (**Fig. 6b**). No significant difference was observed between the WTa2d1xCD38 and rituximab treatments, indicating that WTa2d1xCD38 is as effective as the benchmark treatment in this model. Additionally, treatment with WTa2d1xCD38 significantly prolonged survival compared to the control group, similar to rituximab (**Fig. 6c**). In a second study, we stereotactically engrafted Raji cells into the brains of NSG mice as a model of primary central nervous system lymphoma (PCNSL), an extremely aggressive extranodal B-cell lymphoma with few effective treatment options^28^. We randomized mice to treatment with vehicle control or WTa2d1xCD38. Again, we found that WTa2d1xCD38 inhibited tumor growth and significantly prolonged survival compared to the control group (**Fig. 6e,f**). Together, these findings indicate that WTa2d1xCD38 robustly stimulates anti-tumor responses and could be a highly active therapy for patients with B-cell lymphomas.

**Fig. 6.**
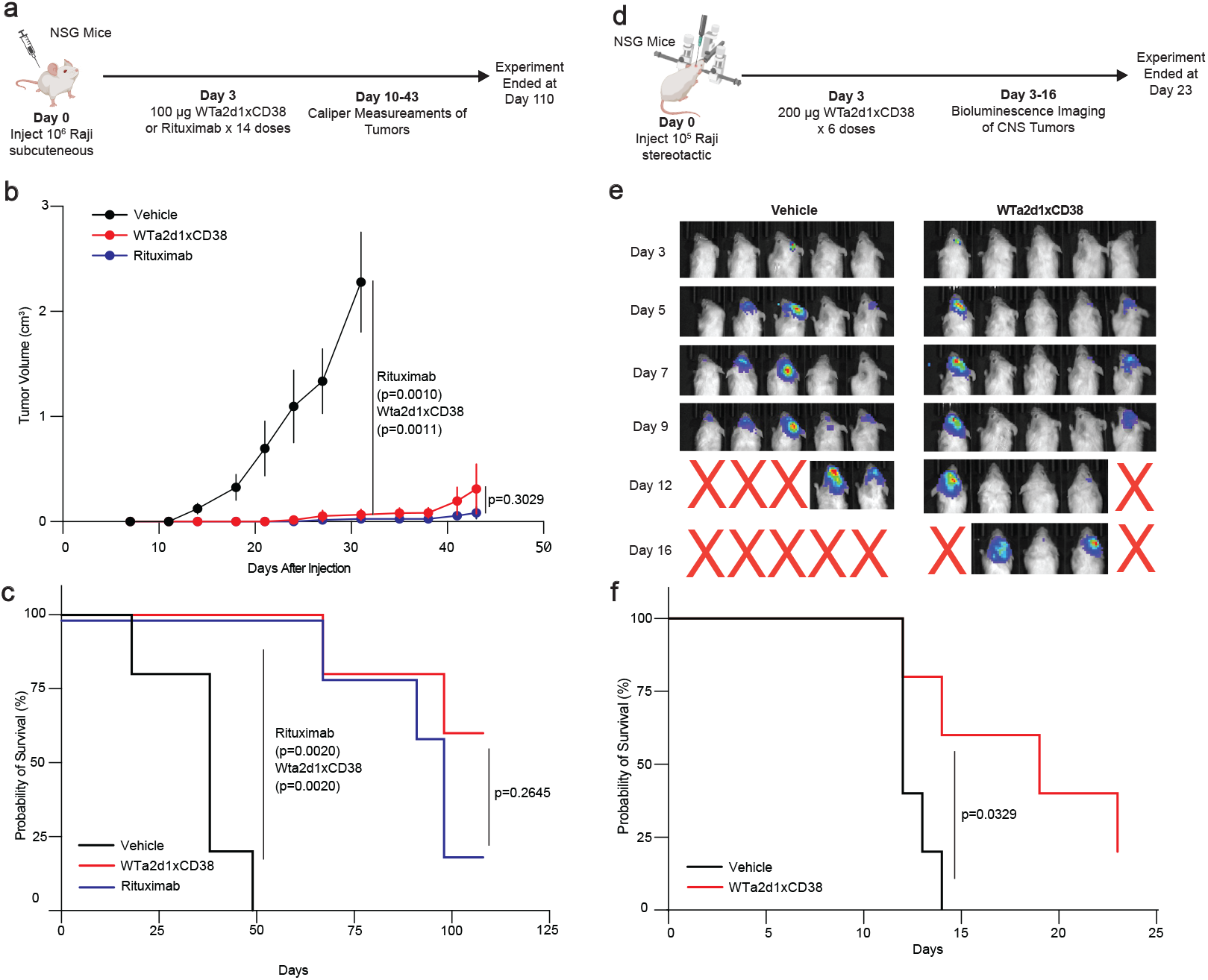
WTa2d1xCD38 exhibits anti-tumor efficacy in xenograft models of aggressive B-cell lymphoma. **a**, Experimental setup of a xenograft experiment using Raji cells engrafted subcutaneously into the flanks of NSG mice. **b**, Growth curves of Raji lymphoma tumors in NSG mice treated with the indicated therapies over time (n = 5 mice per treatment cohort). Tumor growth was evaluated by caliper measurements. Mice were treated with control, 100 µg WTa2d1xCD38, or 100 µg rituximab for 14 days. Data depict mean ± SEM. **c**, Survival analysis of the indicated treatment cohorts. The median survival of Vehicle and Rituximab groups were 38 and 98 days, respectively, whereas the WTa2d1xCD38 was not reached. Log-Rank test was used to evaluate the difference in the probability of survival. Note rituximab curve is minimally nudged for visualization. **d**, Experimental setup of the xenograft CNS lymphoma model experiment using Raji cells engrafted stereotactically into NSG mice brains. **e**, Bioluminescence imaging using luciferase reporters of CNS tumors throughout the experiment period (n = 5 mice per treatment cohort). Mice were treated with vehicle control or 200 µg WTa2d1xCD38 three times per week. **f**, Survival analysis comparing the vehicle control and WTa2d1xCD38 treatment cohorts. Log-Rank test was used to evaluate the difference in the probability of survival between vehicle and treatment conditions.

## Discussion

Monoclonal antibody therapies are among the most successful therapies for B-cell lymphomas and other hematologic malignancies.^29^ These therapies work in part by activating innate immune cell effector functions including macrophage ADCP. This function can be further enhanced by Fc engineering strategies or by blockade of the CD47/SIRPα macrophage immune checkpoint^14,30^. Here, we describe a rational strategy to identify targets of opsonization and antibodies that stimulate macrophage-mediated cytotoxicity of lymphoma cells. Our studies underscore the importance of MHC molecules in protecting lymphoma cells from macrophage attack. Furthermore, our screens identified additional checkpoint molecules, such as CD85j (LILRB1) and CD24, that could be targeted for therapeutic purposes in B-cell lymphoma. Our screens also identified a multitude of unappreciated targets and antibodies that could be further advanced to develop novel therapies that stimulate macrophages to eradicate lymphoma cells. Interestingly, we also identified a number of cell surface targets that were highly expressed on B-cell lymphoma, yet their corresponding antibodies did not stimulate ADCP. These antigens may nonetheless be valuable targets for other therapeutic modalities, such as antibody-drug conjugates, CAR-T cell therapies, or bispecific T-cell engaging therapies.

A limitation of our screens is that they do not sample every possible antigen and every possible antibody that binds to the lymphoma cell surface. However, we provide the first comprehensive surfaceome profiling of B-cell lymphoma that is paired to functional anti-lymphoma responses by macrophages. From our efforts, we have identified consistent principles for macrophage-mediated cytotoxicity across mouse and human studies. Thus, the targets and hits we identified from our screening efforts are true positives that were validated by confirmatory studies in vitro. Furthermore, the results of our screening efforts and combinatorial studies successfully guided and informed the development of a compendium of highly active bispecific antibodies.

Our rapid production and functional evaluation of bsAbs allowed us to create and discover bispecifics that maximally activate macrophage-mediated cytotoxicity of lymphoma cells while minimizing binding to red blood cells. These properties can enhance the therapeutic index for targeting B-cell lymphoma by increasing specificity to the tumor microenvironment. Importantly, bispecifics with the most favorable properties could not be predicted based on their intended targets alone; instead they were identified best by integration across multiple biochemical and functional studies. Thus, this strategy for rapid engineering and evaluation of bispecifics offers opportunities to advance the field of bsAb development. Furthermore, this approach can be applied to other types of cancer or other immune effector cells to yield highly active bispecific antibodies that stimulate immune-mediated cytotoxicity.

Among the most effective bsAbs that we generated was a WTa2d1xCD38 bispecific antibody. This agent exhibited minimal red blood cell binding while robustly stimulating anti-lymphoma responses in vitro and in vivo. CD38 is an established therapeutic target in multiple myeloma, and it is known to be expressed by B-cell lymphomas. However prior trials examining the effects of the anti-CD38 antibody daratumumab observed limited single-agent activity for this drug in B-cell NHL^31^ despite its proven efficacy in multiple myeloma. Our studies suggest that the CD47/SIRPα macrophage immune checkpoint may be a significant barrier that prevents daratumumab from engaging macrophages as immune effector cells. The WTa2d1xCD38 bsAb that we developed may be an ideal way to maximize anti-lymphoma responses by macrophages in this context. Moreover, since macrophages also express CD38, it is possible that this bispecific is acting by multiple functions: (i) opsonizing cancer cells, (ii) blocking the CD47/SIRPα macrophage immune checkpoint, (iii) inhibiting CD38-mediated immunosuppression, and (iv) cross-linking macrophages and cancer cells to enhance immune effector functions **(Fig. 7)**. Indeed, FcγR-blocking experiments showed that while FcγRs are important for the bsAb function, this effect was variable based on the receptor and the cell line, suggesting that the antibody may act through mechanisms in addition to opsonization. Since WTa2d1xCD38 exhibits minimal red blood cell binding, it may be an ideal agent to target macrophage-mediated killing of lymphoma cells while offering a maximal therapeutic window. Moreover, given the expression of CD38 on other hematologic malignancies, this therapeutic may be effective for other disease indications beyond B-cell lymphoma. Our findings indicate these therapeutics hold tremendous potential and should be translated to the clinic for further investigation.

**Fig. 7.**
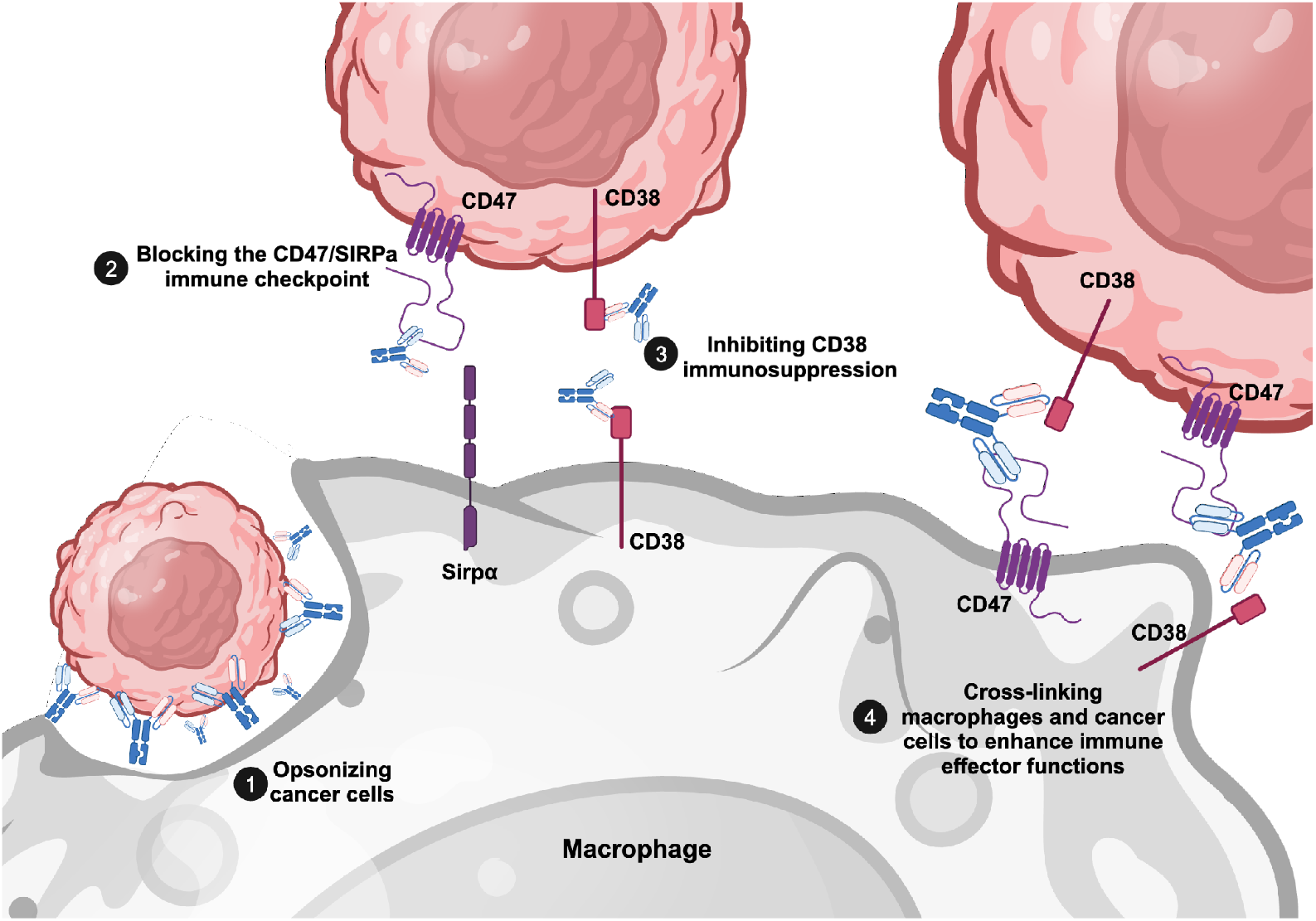
Putative mechanisms of action for the WTa2d1xCD38 bispecific antibody. Representation of the biological functions performed by the bsAb to enhance macrophage-mediated destruction of B-cell lymphoma cells. The WTa2d1xCD38 bispecific can act as an opsonin and engage Fc receptors on macrophages (1). It can also block immunosuppressive pathways on both the cancer and macrophage cell surface (2 and 3). Additionally, it can enhance the biophysical interactions of macrophages and B-cell lymphoma cells, bringing their cell membranes in close proximity to promote phagocytosis (4). Together, these functions maximally activate macrophage-mediated cytotoxicity of aggressive B-cell lymphoma cells.

### Outlook

This study highlights the potential of high-throughput functional screens to identify novel targets and therapies for inducing macrophage-mediated destruction of B-cell lymphoma. The development of a WTa2d1xCD38 bsAb holds promise as a therapeutic for patients since it effectively boosts macrophage anti-lymphoma activity while minimizing the risk of hematologic toxicity. Our findings reveal new opportunities for therapeutic intervention by enhancing the immune system’s innate capabilities, aiming for improved treatment outcomes for patients with aggressive B-cell lymphomas. Overall, our approach provides a blueprint that can be rapidly applied to create new therapies for other types of cancers.

## Methods

### Cell Lines and Culture

A20 (BALB/c B-cell lymphoma) and Raji (human Burkitt lymphoma) cell lines were obtained from ATCC. Toledo (human DLBCL) and SU-DHL-4 (human DLBCL) cell lines were obtained from the Koch Institute High Throughput Services Core. SU-DHL-8 (human large B-cell lymphoma), HBL-1 (AIDS-related NHL), and Daudi (human Burkitt lymphoma) cells were provided by Catherine Wu’s laboratory (Dana-Farber Cancer Institute). All cell lines were cultured in RPMI (Thermo Fisher) supplemented with 10% ultra-low IgG fetal bovine serum (FBS) (Thermo Fisher), 100 units/mL penicillin, 100 µg/mL streptomycin, and 292 µg/mL L-glutamine (Thermo Fisher). Cell lines were maintained in humidified incubators at 37 °C with 5 % carbon dioxide.

MHC-I knockout A20 and Raji variants were made by CRISPR-Cas9 genome editing using a Cas9 endonuclease. Mouse TAP1 gRNA (GCGGCACCTCGGGAACCAACAGG), human B2M gRNA (CGTGAGTAAACCTGAATCTTTGG), Cas9 endonuclease and universal tracrRNA were obtained from Integrated DNA Technologies (IDT). The crRNA:tracrRNA complex was ensembled following the manufacturer’s protocol and delivered by electroporation using the Neon Transfection System (Thermo Fisher).

### Generation of fluorescent cell lines

StayGold+ cell lines were generated by transduction of cells with a lentivirus encoding hStayGold (Genbank LC593679.1) under the control of an EF1-ɑ promoter (Vectorbuilder) followed by selection with puromycin. GFP+ cell lines were generated by lentiviral transduction of cells using CMV-GFP-T2A-Luciferase pre-packaged virus (Systems Bio). Transduced cells were then sorted for stable GFP expression. To generate a stable mScarlet Toledo line, a DNA fragment encoding the monomeric mScarlet red fluorescent protein was synthesized (IDT) and cloned into an AAVS1-homology donor vector previously published^32^ using HiFi DNA Assembly (NEB) following restriction with XbaI and MluI digest of the targeting donor vector and inclusion of homology arms on the synthesized mScarlet fragment. The AAVS1-mScarlet donor template vector was introduced in the cells along with an AAVS1-targeting Cas9 PX458 (#48138-Addgene) plasmid containing the sgRNA sequence: GGGGCCACTAGGGACAGGAT using nucleofection according to the manufacturer’s specifications (Lonza 4D Nucleofector, B-cell protocol) at a DNA mass ratio of 4:1 donor to Cas9 plasmid. The cells were selected using puromycin driven by the endogenous AAVS1 transcription via a splice acceptor and 2A chysel sequence upstream of the puromycin cassette in the AAVS1-homology donor construct. Cells were then expanded and further enriched for reporter expression using fluorescence-assisted cell sorting. Cells were frozen immediately after FACS enrichment, banked, and thawed for use in experiments.

### Mouse macrophage derivation

Primary murine macrophages were derived from syngeneic BALB/c mice as previously described^18^. In brief, long bones from mice were collected and bone marrow was mechanically extracted using a mortar and pestle. Cells were washed with PBS then subjected to ACK lysis (ThermoFisher) to deplete red blood cells. Unfractionated bone marrow cells were plated on Petri dishes (Corning) with 20 ng/mL murine M-CSF (Peprotech) for at least 7 days to differentiate into macrophages. Macrophages were washed, removed from plates using TrypLE (Invitrogen) and cell lifting, and used for experiments or replated as necessary. Macrophages were generally used for experiments between days 7-21 of culture.

### Human macrophage derivation

Primary human macrophages were derived from peripheral blood monocytes of anonymized healthy human blood donors as previously described^18,33^. Briefly, leukocyte reduction collars from anonymized blood donors were obtained from the Crimson Core Biobank (Brigham and Women’s Hospital). Monocytes were isolated using StraightFrom Whole Blood CD14+ microbeads (Miltenyi) using an AutoMACS Pro or AutoMACS Neo Separator (Miltenyi). Purified monocytes were cultured in IMDM (ThermoFisher) + 10% low IgG FBS (ThermoFisher) + 1x penicillin, streptomycin, and glutamine (ThermoFisher) containing 20 ng/mL human M-CSF (Peprotech) for 7 days to differentiate into macrophages. Macrophages were generally used for experimentation between days 7-21 of culture and removed from plates by trypsinization and cell lifting before replating as necessary.

### Long-term co-culture assays

Long-term co-culture of murine or human macrophages and lymphoma cells was performed using fluorescently labeled target cells as previously described^18^. Phenol-red free IMDM (Thermo Fisher) supplemented with 10% ultra-low IgG fetal bovine serum, 100 units/mL penicillin, 10 μg/mL streptomycin, and 292 µg/mL L-glutamine, and 20 ng/mL M-CSF was used. For antibody library screens, Lyoplate Mouse and Lyoplate Human Cell Surface Marker Screening Panel plates (BD) were used. Library plates containing lyophilized antibodies were centrifuged at 300 x g for 5 minutes before reconstitution with 140 μl of IMDM using an ASSIST PLUS pipetting robot (Integra). To set up co-cultures, 20 μl containing 1×10^4^ macrophages and 2×10^3^ target cells (4:1 E:T ratio) were added to each well of a 384-well plate using an ASSIST Plus (Integra). Subsequently, 20 μl from each Lyoplate well was added to achieve a final working concentration of 6.55 μg/ml. Finally, an additional 20 μl medium alone or medium containing anti-CD47 or anti-CD20 was added to the wells. This achieved a final working concentration of 10 μg/mL of anti-CD47 or anti-CD20 as appropriate. Cells were then co-cultured for 156 h (∼6.5 days), with whole-well imaging of phase contrast and green and red channels performed every 8 hours using an Incucyte S3 system. Automated imaging analysis was performed using Incucyte Analysis Software.

### FACS analysis of library antibodies binding

To evaluate specific binding interactions, 1×10^5^ target cells were added in 100 μl to each well of the Lyoplate plate containing 20 μl of purified antibody. The plate was then incubated on ice for 30 minutes to facilitate binding. For murine binding, biotinylated goat anti-mouse, anti-rat, and anti-Syrian hamster antibodies, at a concentration of 1.25 μg/mL each, were added at 100 μl per well, while biotinylated anti-Armenian hamster antibody, at a concentration of 0.6 μg/mL, was added at the same volume, to each corresponding well. The plate was then incubated for an additional 30 minutes on ice. After washing, plates were incubated with Alexa Fluor 647 streptavidin, diluted 1:4000, to achieve a final concentration of 0.5 μg/mL for 30 minutes on ice, followed by another round of washing. For human binding, Alexa Fluor 647 Goat anti-mouse Ig and Goat anti-Rat Ig (BD) were directly added as the first step in each according well (dilution 1:200), following the same incubation times. Cells were then resuspended with 100 μl of FACS Staining Buffer (BD) for flow cytometry analysis, performed using a BD LSR Fortessa flow cytometer. The mean fluorescence intensity (MFI) of the bound streptavidin-labeled antibodies was measured to quantify the extent of binding. Data analysis was conducted using FlowJo (version 10.9), and the MFI was calculated accordingly.

### Scfv Library and Plasmid Generation

Antibody sequences were obtained from publicly available sources and were directed against the following targets: CD20 (rituximab, IMGT 161), CD85 (US Patent Application Pub. No.: 20230235055A1), CD71 (delpacibart, IMGT 1374), CD38 (daratumumab, IMGT 301), CD40 (dacetuzumab, IMGT 232), CD47(CV1^14^), CD47(WTa2d1^14^), CD24-1^34^, CD24-2 (US Patent Application Pub. No.: 20210213055A1), CD184 (ulocuplumab, IMGT 483), PD1 (nivolumab, KEGG D10316). The sequences were numbered using the Kabat numbering system and their VH and VL regions were fused through a (GGGGS)X3 linker. The sequence was reverse-translated and optimized for the *Homo sapiens* codon usage table. Gene blocks ordered from Twist Biosciences containing these sequences were engineered with flaking 5’ and 3’ multiple cloning sites containing EcoRI/BamHI and NotI. The cloning was carried over using restriction enzyme digestion followed by T4 DNA ligation and bacterial transformation. Individual colonies were subjected to Nanopore sequencing (Quintara Biosciences) to verify correct gene insertion. The ZymoPURE™ Plasmid Miniprep Kit (endotoxin levels ≤ 1 EU/µg of plasmid DNA) was used to extract plasmid DNA from transformed bacteria, which was subsequently used for transfection.

### Expression and Purification of Recombinant Antibodies

Purified pairs of IgG1-Knob and IgG1-Hole plasmids were co-transfected to a final concentration of 1 µg/mL of each into 900 μl of Expi293F cells at 3×10^6^ cells/mL (Thermo Fisher) in 96 deep-well plate (USA Scientific) with ExpiFectamine 293 Transfection Kit (Thermo Fisher Scientific) following the manufacturer’s recommendation and incubated at 37°C, 8% CO2 with shaking at 900 rpm for 7 days. Cells were pelleted at 3,000 xg for 20 min and the antibody supernatants were diluted 2-fold with PBS and used for further analysis.

### ELISA

Immulon 4HBX Flat Bottom plates (Thermo Fisher) were coated with 1:1000 dilution in PBS of anti-human IgG Fc (Jackson Immunoresearch) overnight at 4 °C. The following day, the plate was washed three times with PBST and blocked with 1% BSA in PBST for one hour at room temperature. The plate was then washed with PBST, loaded with the diluted transformed supernatants from Expi293F cells, and incubated for another hour at room temperature. Upon washing again, anti-human IgG H+L (Jackson Immunoresearch) was added to the plate at 1:100,000 dilution in 2% (w/v) non-fat dry milk in PBST and incubated for another hour at room temperature. Finally, another set of washes was performed and 1-Step Turbo TMB Elisa Substrate (ThermoFisher) was added to the plate and the reaction was quenched with 10% phosphoric acid after 3 min. The absorbance was assessed at 450 nm.

### Phagocytosis assays

In vitro phagocytosis assays were performed as previously described^14^. Briefly, CFSE+ Raji and mScarlet+ Toledo cells were used as target cells. Labeling of Raji cells with CFSE (Thermo Fisher) was performed according to the manufacturer’s instructions. Live cells were collected, washed, and co-cultured with murine macrophages at a target macrophage ratio of 4:1. Cells were co-cultured in the presence or absence of different antibodies at a final concentration of 10 µg/mL. Cells were co-cultured for 2 hours in serum-free IMDM in round-bottom ultra-low attachment 96-well plates (Corning). After the incubation period, cells were washed and analyzed by flow cytometry. Macrophages were identified using APC anti-SIRPα (BioLegend) and target cells were identified by CFSE fluorescence. Phagocytosis was quantified as the percentage of macrophages that contained CFSE signal. Phagocytosis was normalized to the maximal response. Dose-response curves were generated using Prism version 9.2.0 (GraphPad).

### Recombinant protein production and purification

WTa2d1xCD38 plasmids were produced using ZymoPure MaxiPrep kit following manufacturer instructions. Plasmid concentration and purity were assessed using Nanodrop (Thermo Fisher) and 1 µg/mL of each was employed to transient transfect Expi293F cells (Thermo Fisher) following manufacturer instructions. Subsequently, the culture was pelleted at 15,000 xg for 15 min and the supernatant was loaded in a Protein A column (Cytiva) equilibrated with TBS (pH 7.2). The column was washed with 25 ml of TBS and the protein was eluted with 100mM Glycine and 100mM NaCl solution (pH 3.0). The eluate was brought to physiological pH with 1M Tris (pH 8.0) and the solution was buffer exchanged to PBS using either dialysis bags (ThermoFisher) or spin-columns (Cytiva). Protein concentration was obtained using a Nanodrop (Thermo Fisher by taking the absorbance at 280 nm.

### Xenograft mouse models

NSG mice were obtained from Jackson labs (Strain: #005557) and used for experiments when approximately 6-12 weeks of age. Age and sex-matched mice were engrafted subcutaneously with 1×10^6^ GFP+ Raji cells in 50% (v/v) Matrigel Matrix (Corning) and PBS. Mice were randomized to treatment cohorts and then subjected to intraperitoneal treatment in vivo with vehicle control, 100 µg WTa2d1xCD38, or 100 µg rituximab biosimilar (BioXCell). Mice were treated 5-7 times per week for 14 days. Tumor dimensions were measured by caliper twice per week and used to calculate tumor volumes using an ellipsoid formula: length x width x width x π/6.

Eight-week-old female NSG mice (Charles River Labs, Strain: #614) were used to develop a model of primary central nervous system lymphoma (PCNSL). Mice were initially anesthetized with isoflurane (1-2%), followed by intradermal injection of ketamine (80-100 mg/kg) and xylazine (5-10 mg/kg) for anesthesia maintenance and analgesia. A total of 10^5^ Raji cells in 5 µl of PBS were injected intracerebrally using a Hamilton syringe with a 26-gauge needle at a rate of 1 µl/min, guided by stereotactic coordinates (1 mm anterior, 1.8 mm lateral right to the bregma, and 2.5 mm deep from the dura) on a Stoelting Just For Mice platform. Mice were then randomized into treatment groups and received intravenous injections of either vehicle control or 200 µg of WTa2d1xCD38 starting on day 3 post tumor injection, administered three times per week for a total of six injections. Tumor growth was monitored by bioluminescence imaging (BLI) using an IVIS Spectrum system (PerkinElmer) twice a week starting on day 3 post-intracerebral injection. Tumoral size was analyzed and quantified using Living Image software (PerkinElmer), and the total photons per second (ph/s) were recorded.

For survival analysis, mice were euthanized when humane experimental endpoint criteria were met, including neurological symptoms (seizures, circling or hind limb paralysis) or a significant weight loss (>20%).

### Statistical Analysis

GraphPad Prism (v10.2) was used to perform correlation and grouped analysis. We performed a Pearson correlation test and a linear regression to evaluate the relationship between the functional and biochemical properties of the antibodies. A two-way ANOVA with correction for multiple comparisons at the indicated time point was used to detect the differences between different sample groups in both in vivo (tumor measurement) and in vitro datasets. If necessary, Tukey’s multiple comparisons test was performed to distinguish the differences within groups. For paired group analysis, a Student two-tailed paired T-test was performed to elucidate the differences in grouped data. Survival analysis was performed using the log-rank (Mantel-Cox) test. Significant differences were determined as p<0.05. For the long-term co-culture and in vivo experiments, the results are expressed as mean ±SEM, whereas for the remaining experiments, the results are expressed as mean ±SD.

For other statistical analyses, Python (v.3.10) was used. The datasets were normalized using sklearn MinMaxScaler function and Seaborn was used to calculate Euclidean distance and perform hierarchical clustering of the different therapeutic combinations in the human system. For K-means clustering, the dataset was first normalized using sklearn StandardScaler (z-score normalization) and then fitted using skelarn’s K-means clustering function. The silhouette method was used to identify the optimal number of clusters.

### Reporting Summary

Further information on research design is available in the Nature Research Reporting Summary linked to this article.

## Data Availability

The main data supporting the results in this study are available within the paper and its Supplementary Information. Source data for the figures will be provided with this paper.

## Acknowledgements

We thank members of the Weiskopf lab as well as members of the Tobiloba Oni and Robert Weinberg labs for experimental assistance, reagents, and insightful discussions. We thank Ferenc Reinhardt, Donna Hicks, Elinor Eaton, Joana Liu Donaher, Nicholas Polizzi, and Beverly Dobson for support. We thank Patrick Autissier and the Whitehead Institute Flow Cytometry Core Facility and the Bioinformatics and Research Computing Core Facility. Experimental diagrams were created using Biorender.

## Funding

KW reports support from the Valhalla Foundation, AACR-AstraZeneca Career Development Award for Physician-Scientists in Honor of José Baselga, an anonymous grant, Richard Reisman in honor of Jane Reisman, Department of Defense [W81XWH2210141], National Cancer Institute [R01 CA279259], and the Research Foundation for the Treatment of Ovarian Cancer. CPG reports support from a EACR Travel Fellowship. MC was supported by research funding from Asociación Española Contra el Cáncer (AECC) [LABAE18014CRES] and from Instituto de Salud Carlos III, Fondo de Investigaciones Sanitarias [PI21/01190], co-sponsored by the European Union FEDER program “Una manera de hacer Europa”. The content of this manuscript is solely the responsibility of the authors and does not necessarily represent the official views of the NIH, fundació BBVA and Cellex, or other funding agencies.

## Author contributions

CPG, JR, JV, AM, JB, KV, TW, and KW designed and performed in vitro assays. JR, KW and CPG designed experiments and analyzed data pertaining to bispecific antibodies. JR, AM, KW, CPG and PFG designed and performed mouse experiments. MC and KW performed experimental oversight and funding acquisition. CH generated cell lines. CPG, JR, and KW wrote the first draft of the manuscript. All authors reviewed and edited the manuscript and are responsible for its contents.

## Competing interests

JR, CPG, KV, AM, and KW have filed US patent applications related to this work. KV is a former employee and equity owner of DEM Biopharma. MC has received research funding from Janssen, Genentech and AstraZeneca. KW reports patents/royalties (Stanford University, Whitehead Institute, Forty Seven, Gilead Sciences, ALX Oncology, DEM Biopharma); co-founder, scientific advisory board member, and equity holder (ALX Oncology, DEM Biopharma, Solu Therapeutics), stock ownership (Ginkgo Bioworks). The other authors declare no competing interests.

---

**Supplementary Fig. 1.**
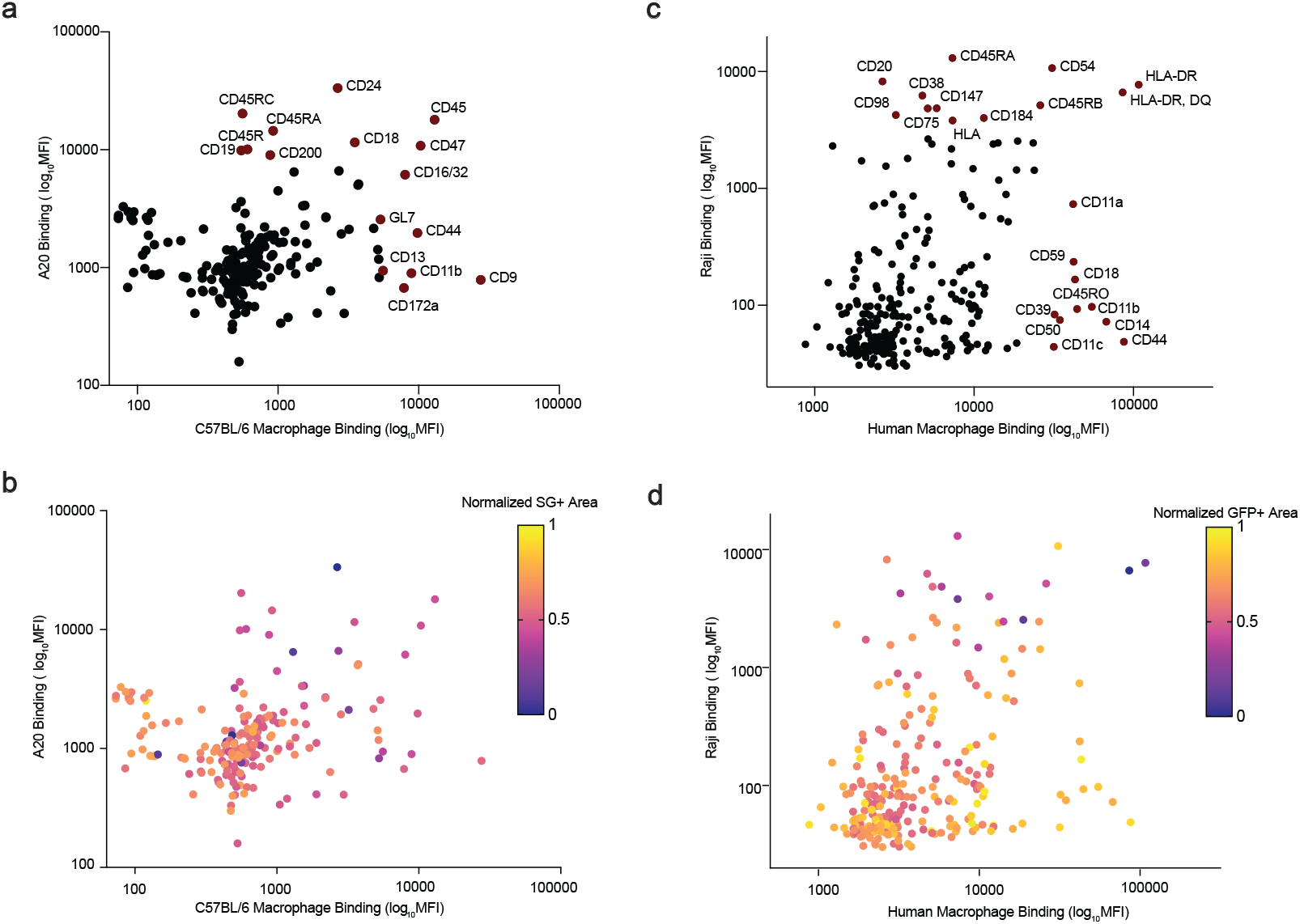
Binding of antibodies from libraries to B-cell lymphoma cells and macrophages. **a**, Scatter plot depicting cell-based binding of each individual antibody from the murine antibody library binding to A20 lymphoma cells and C57BL/6J macrophages. Mean fluorescence intensity (MFI) was assessed by flow cytometry using Alexa 647-conjugated streptavidin for detection. **b**, Scatter plot depicting results of cell-based binding of each individual antibody from the human antibody library to Raji lymphoma cells and primary human macrophages combined from n = 3 independent donors. Binding was detected using Alexa 647-conjugated anti-Ig secondary antibodies. **a,b**, The target antigens highlighted exceeded the 95^th^ percentile for binding to either cancer or macrophages cells. **c,d**, Multiple variable scatter plot depicting the relationship between the binding of the antibody library to lymphoma cells and/or macrophages and the functional anti-tumor effects of each antibody in co-culture with macrophages (color scale) in the murine (**c**) and human (**d**) systems. Color scale represents the cancer cell area at the last time point in co-culture assays, with blue indicating greater anti-lymphoma activity and yellow indicating lesser anti-lymphoma activity.

**Supplementary Fig. 2.**
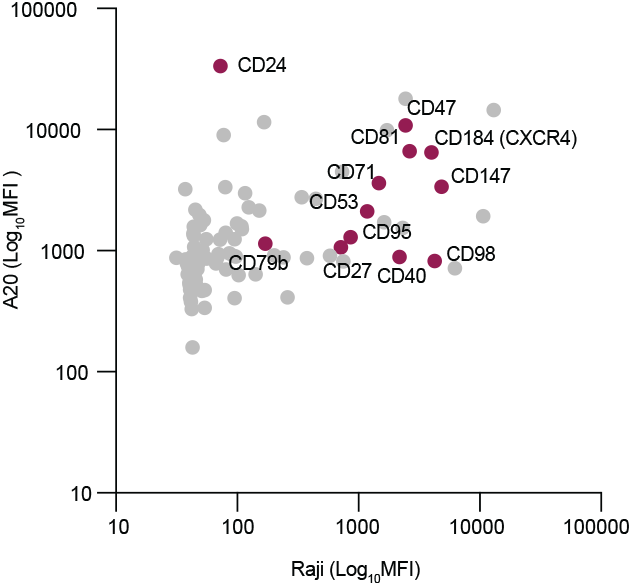
Comparison of antigens and antibodies evaluated from libraries in the murine and human systems. Scatter plot depicting the relationship between binding of 92 shared targets in both human (Raji) and murine systems (A20). The highlighted antigens are the selected antibody hits that exceeded the 95^th^ percentile of functional activity from the co-culture assays with macrophages and B-cell lymphomas as single agents. Binding (MFI) was determined by flow cytometry using Alexa 647-conjugated streptavidin (A20) or Alexa 647-conjugated anti-Ig antibodies (Raji) for detection.

